# Symmetric arrangement of mitochondria:plasma membrane contacts between adjacent photoreceptor cells regulated by Opa1

**DOI:** 10.1101/635094

**Authors:** Ingrid P. Meschede, Miguel C. Seabra, Clare E. Futter, Marcela Votruba, Michael E. Cheetham, Thomas Burgoyne

**Affiliations:** UCL Institute of Ophthalmology, London, EC1V 9EL; CEDOC, NOVA Universidade Nova de Lisboa, Lisbon, Portugal; School of Optometry and Vision Sciences, Cardiff University, Cardiff, UK; Cardiff Eye Unit, University Hospital Wales, Cardiff, UK

**Keywords:** rod photoreceptors, mitochondria, contact sites, tethers

## Abstract

Mitochondria are known to play an essential role in photoreceptor function and wellbeing that enables normal healthy vision. Within photoreceptors they are elongated and extend most of the length inner segment, where they supply energy for protein synthesis and the phototransduction machinery in the outer segment as well as acting as a calcium store. Here we examined the arrangement of the mitochondria within the inner segment in detail using 3D electron microscopy techniques and show they are tethered to the plasma membrane in a highly specialised arrangement. This includes mitochondria running alongside each other in neighbouring inner segments, with evidence of alignment of the cristae openings. As the pathway by which photoreceptors meet their high energy demands is not fully understood, we propose this to be a mechanism to share metabolites and assist in maintaining homeostasis across the photoreceptor cell layer. In the extracellular space between photoreceptors, Müller glial processes were identified. Due to the often close proximity to the inner segment mitochondria, they may too play a role in the inner segment mitochondrial arrangement as well as metabolite shuttling. OPA1 is an important factor in mitochondrial homeostasis, including cristae remodelling; therefore, we examined the photoreceptors of a heterozygous *Opa1* knock-out mouse model. The cristae structure in the *Opa1*^+/−^ photoreceptors was not greatly affected, but there were morphological abnormalities and a reduction in mitochondria in contact with the inner segment plasma membrane. This indicates the importance of key regulators in maintaining this specialised photoreceptor mitochondrial arrangement.

## Introduction

Vertebrate photoreceptors are specialised neurons that provide vision by transducing light into electrical signals. The combination of phototransduction, neurotransmitter utilization, protein synthesis and transport and repolarisation after depolarisation makes the energy consumption of photoreceptors greater than all other cell types in the body (1, 2). As a consequence, failure to fulfil their energy requirements often results in visual problems including blindness. These include a number of diseases such as Leber’s Hereditary Optic Neuropathy, Dominant Optic Atrophy, and Leigh syndrome that lead to malformed or dysfunctional mitochondria (3, 4). Most of the mitochondria of photoreceptors are housed within the inner segment (IS) region and are typically elongated running along the long axis of the photoreceptor. Within the IS mitochondria are well situated for uptake of extracellular metabolites via channels on the IS plasma membrane (PM) and can provide the necessary energy for protein synthesis and for the phototransduction machinery of the adjoined outer segment (OS). Furthermore, photoreceptor mitochondria have been shown to act as a calcium store (5, 6). Ca2+ regulation is crucial for signalling including phototransduction, membrane excitability, energy metabolism, cytoskeletal dynamics, and transmitter release (7–10).

Dominant optic atrophy (DOA) is an autosomal disease that affects the optic nerves, leading to reduced visual acuity and preadolescent blindness (11). The most common cause of DOA are mutations in *OPA1* that codes for a dynamin-related guanosine triphosphatase (12, 13). OPA1 is required for lipid mixing and fusion of the mitochondrial inner membranes (14). In addition to the optic nerve, *OPA1* has been shown to be expressed in the retina in the photoreceptor IS (15). The role and impact of its loss of function in photoreceptors has not been well studied.

When examining published transmission electron microscopy (TEM) images of mouse photoreceptors, the tissue is usually orientated longitudinally. When viewed like this, it is difficult to determine the fine positioning of mitochondria, and how this relates to the energy and storage demands within the photoreceptor IS. In this study, we set out to examine the mitochondria arrangement in detail throughout the depth of entire photoreceptor IS, as well as the cristae architecture using 3D electron microscopy analyses. As a consequence, we discover new insight into the arrangement, and morphology of mitochondria within the IS, which includes the first description of extensive contact sites between mitochondria and the PM in mammalian cells. This sheds light on the importance and myriad of roles that mitochondria play in photoreceptors and how this can be affected in disease models, such as we observed in the heterozygous *Opa1* knock-out (KO) mouse model.

## Results

### Mitochondria from neighbouring photoreceptor ISs are aligned to run side-by-side

To study the 3D arrangement of mitochondria within photoreceptor IS, P20 wild-type mouse eyes were prepared for serial block face scanning electron microscopy (SBFSEM). Single images from the SBFSEM data showed the mitochondria to be positioned in close proximity to the PM throughout the entire IS (Fig. 1 A). The arrangement of mitochondria varied at different depths of the IS. Within most of the IS, mitochondria appeared to cluster adjacent to mitochondria in neighbouring photoreceptors. Reduced clustering was observed at the tip of the IS, close to the OS (distal end of the IS) or the Golgi (proximal end of the IS). By modelling a portion of SBFSEM data, the mitochondria from neighbouring photoreceptors can be seen to clearly run alongside each other through most of the depth of the IS (Fig. 1 B and Fig. S2). Some mitochondria were seen to be forked shaped and run alongside multiple mitochondria from neighbouring cells (as shown by mitochondria modelled in blue in Fig. 1 B). The mitochondrial arrangement determined from the SBFSEM data correlated with observations made from conventional TEM samples of 6 month old and P20 wild-type mice (Fig. 1 C, D). Mitochondria from neighbouring photoreceptors were seen running side-by-side in longitudinal samples as well as arranged in doublets or triplets in transversely orientated samples. The same mitochondrial arrangement was observed in rod and neighbouring cone photoreceptors (Fig. S1) that were identified by the higher density of less electron dense mitochondria (16). In the extracellular space between photoreceptor IS, often positioned between the mitochondria, small circular membranous structures were observed within TEM images (white arrowheads in Fig. 1 D, E). When examined in the SBFSEM data these were found to be projections that run up between the IS (Fig. S2).

**Fig. 1.**
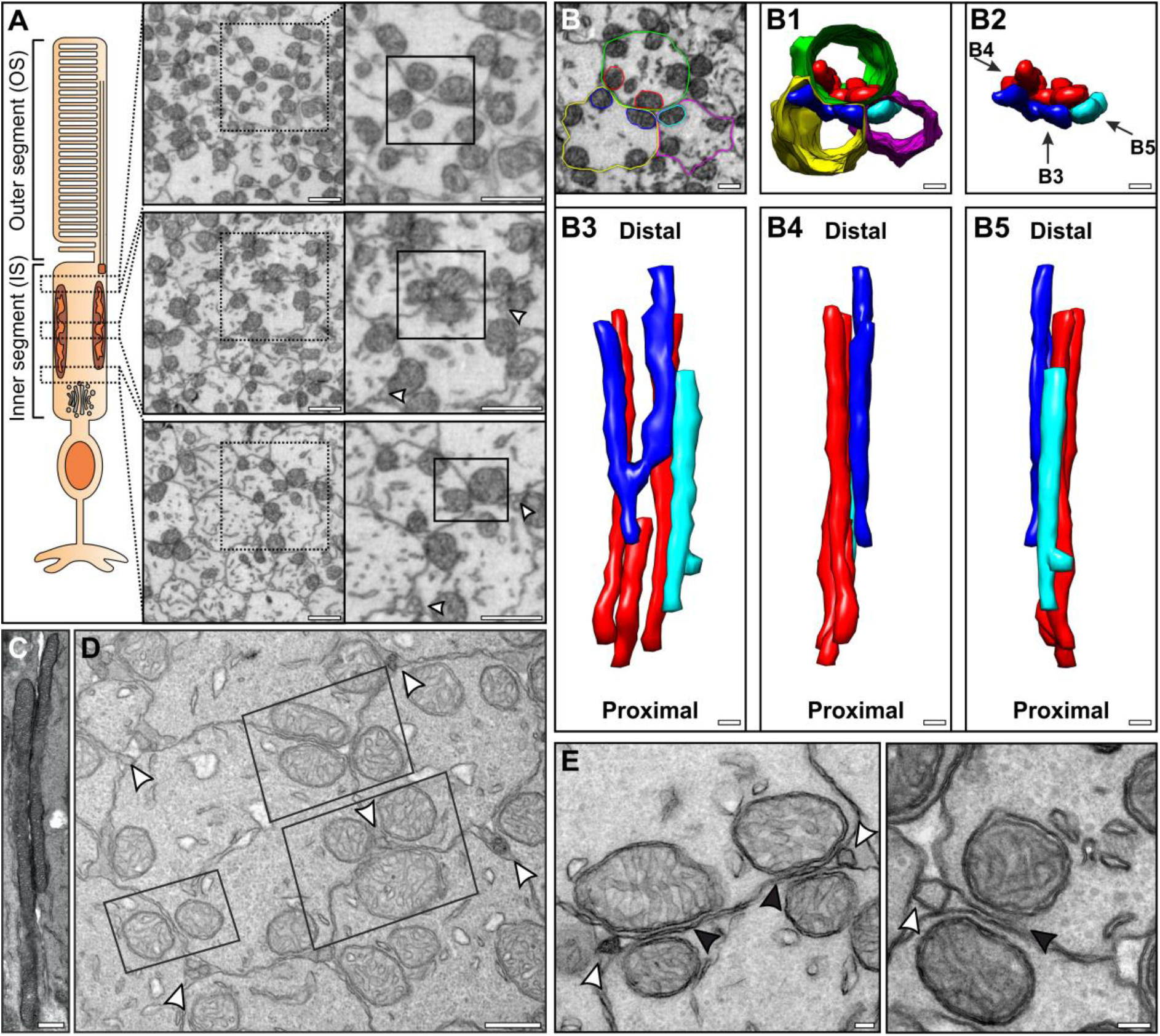
Mitochondria from neighbouring photoreceptors align through the depth of the IS with projections visible nearby in the extracellular space. (A) SBFSEM images of mouse photoreceptors cross sections where there is apparent alignment of mitochondria from neighbouring photoreceptors that is most prominent in the mid region of the IS as shown by the box with a solid outline. (B) Segmentation and modelling of select mitochondria and PMs from three ISs, showing the mitochondria running alongside each other. (C) Longitudinal TEM section showing mitochondria from neighbouring ISs running side-by-side. (D) TEM cross section showing mitochondria arranged in pairs or triplets between neighbouring ISs highlighted by the boxed region. Membrane projections were seen in between the ISs in close proximity to the mitochondria (white arrowheads). These are shown at a higher magnification in (E) by the white arrowheads as well as alignment of neighbouring IS mitochondria indicated by the black arrowheads. Scale: (A) 1μm, (B-D) 500nm and (E) 100nm.

### Mitochondria are tethered to the PM and the cristae between mitochondria from neighbouring cells appear to be aligned

By examining the IS mitochondria at high magnification within TEM images, electron dense tethers were detected between the mitochondrial outer membrane and the PM (Fig. 2 A). In addition, a high degree of consistency was found when measuring the distance between the two membranes (Fig. 2 B), which was found to be 10.77 nm (±0.28). To examine the tethering and mitochondria structure in 3D at a higher resolution than is achievable by SBFSEM, tomograms were generated (Fig. 2 C). Within the tomographic data the tethers were resolved and could be seen bridging the outer mitochondrial and PMs (Fig. 2 D and Movie S1, S2). When examining the mitochondria cristae, at different depths within the tomograms, the opening of the cristae appeared to be aligned (77.84% ± 0.98% SE) between neighbouring mitochondria (Fig. 2 E). This was further confirmed by modelling the mitochondria membranes indicating many of the cristae openings are opposed to each other (black dotted line in Fig. 2 F & Movie S1 – S3).

**Fig. 2.**
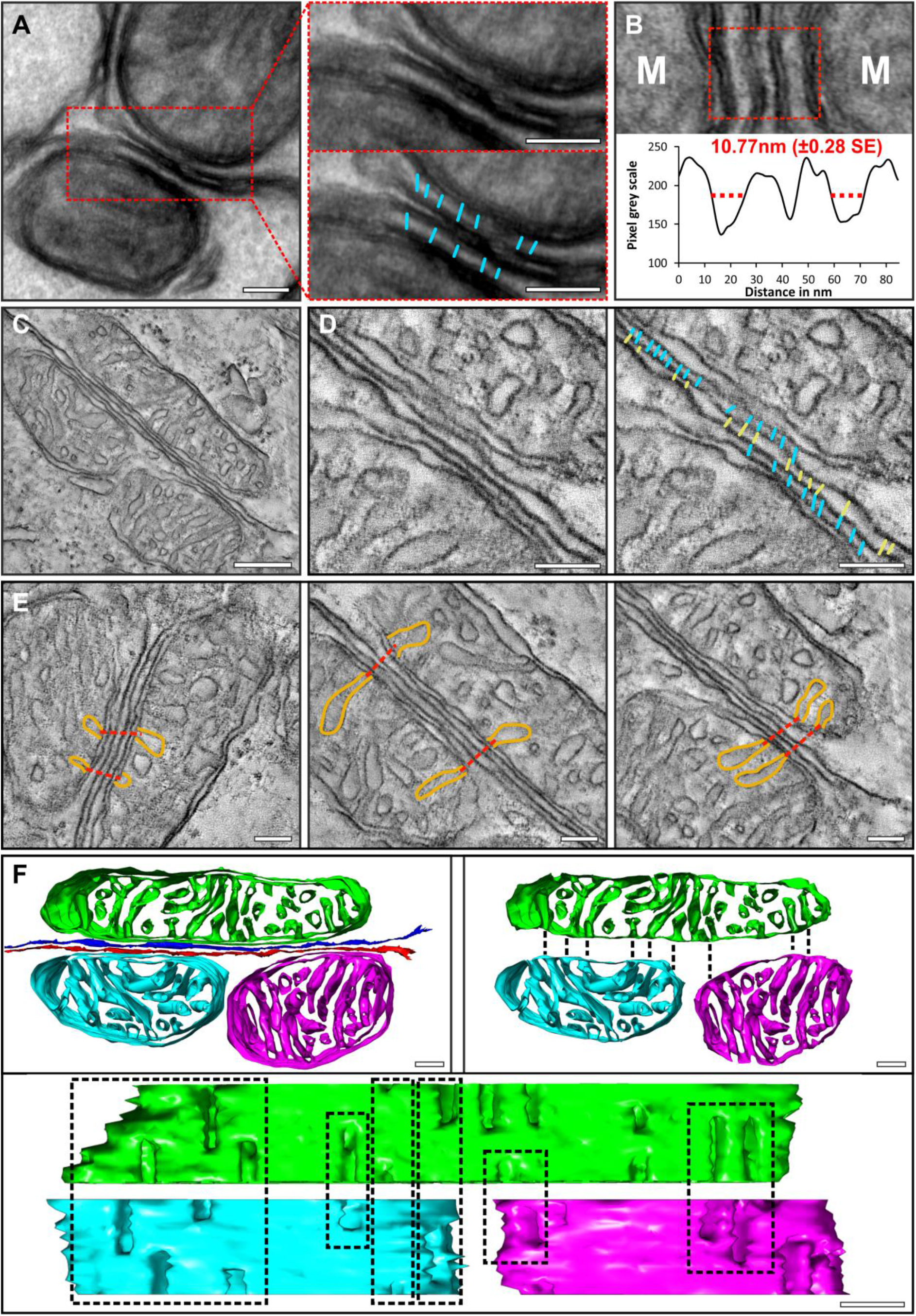
Visible tethers connect mitochondria to the IS PM and there is alignment of the mitochondrial cristae between neighbouring cells. (A) Tethers between the mitochondrial outer membrane and the PM are visible in a thick (200nm) TEM section as highlighted in cyan. (B) The distance between the mitochondrial outer member and PM is well conserved at 10.77nm (± 0.28 SE). (C-F) Tomography reconstructions and resulting models of IS mitochondria. (C) Is a single slice from a tomogram reconstruction and when focusing on the PM (D) tethers can be seen connecting the mitochondria to the PM (cyan) as well as structure detected between the two ISs (yellow). (E) Single slices from the tomograms showing alignment of cristae openings of mitochondria from neighbouring photoreceptors. (F) A model of the mitochondria generated from a tomogram reconstruction. When removing the outer membrane from the model, there is visible alignment of the cristae openings. Scale: (A) 100nm, (C) 250nm, (D-F) 100nm.

### Neighbouring cell mitochondria cristae alignment was not observed within the retinal pigment epithelium

To investigate if the alignment of the cristae across mitochondria of neighbouring cells is a general phenomenon in the retina, the retinal pigment epithelium (RPE) was examined. At the RPE lateral membrane, mitochondria are closely associated to the PM (Fig. 3 A, distance between mitochondrial outer membrane and PM measured as 10.45nm ± 0.53nm SE), similar to what we observed in the photoreceptor IS. Most of these were not found to be positioned adjacent to mitochondria of neighbouring cells (42.01% ± 5.25% SE) the ones that were had little or no cristae alignment (Fig. 3 B). This was in contrast to the photoreceptor IS cristae that were found to be aligned in both longitudinally and transversely orientated tissue samples (Fig. 3 C, D).

**Fig. 3.**
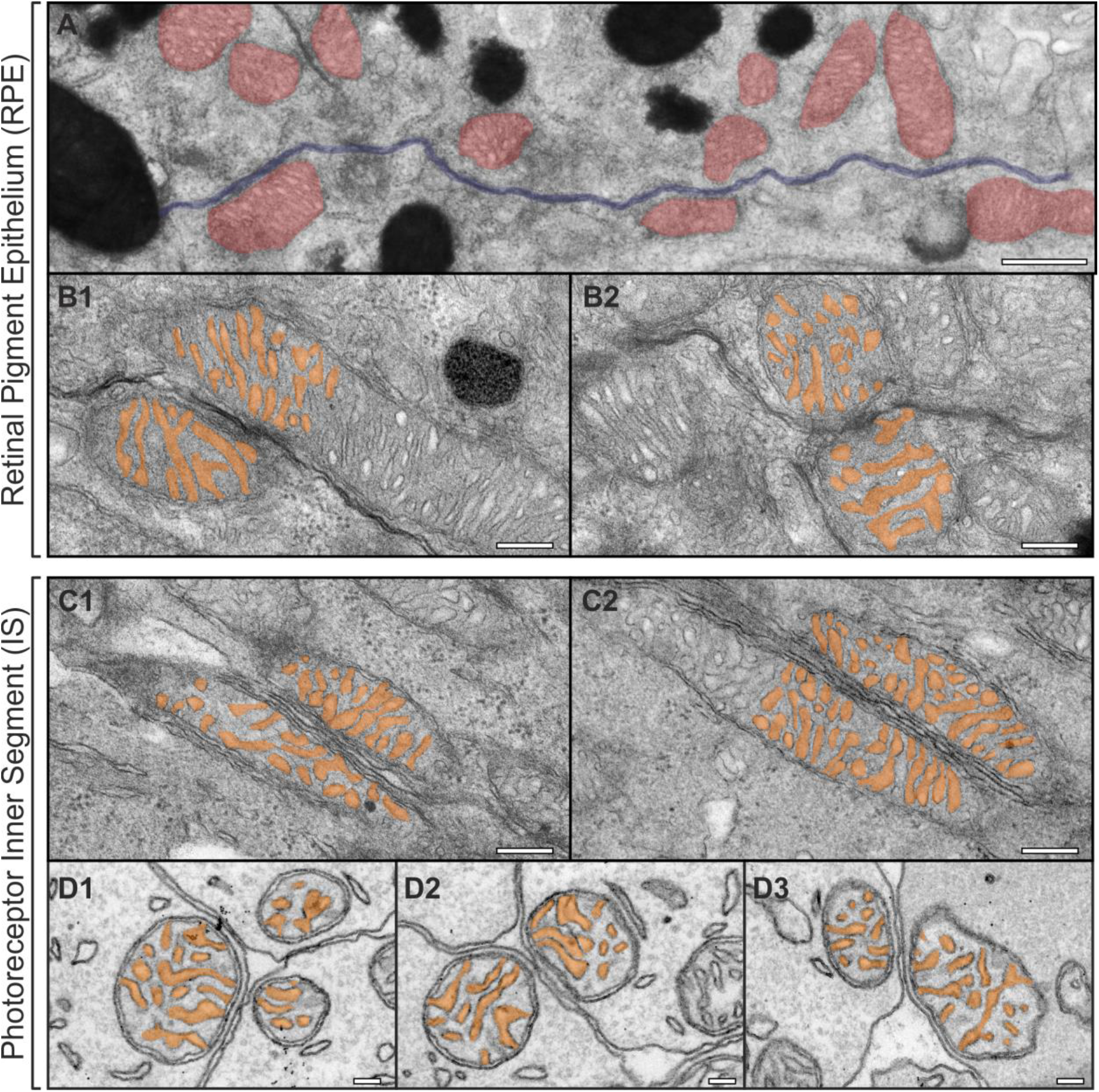
The mitochondrial alignment is not observed in the RPE when compared to photoreceptor ISs. (A) At the lateral RPE border mitochondria (in red) are in contact with the PM (in blue). (B) TEM image of mitochondria from neighbouring RPE cells including false coloured cristae show little alignment of the cristae openings. Whereas the cristae opening alignment is seen in (C) longitudinally orientated and (D) Transverse TEM sections of neighbouring IS mitochondria. Scale: (A) 500nm, (B-C) 200nm and (D) 100nm.

### Müller Glial processes run between photoreceptor ISs

The small circular membrane observed in transversely orientated mouse retina (Fig. 4 A) were seen as tubular projections running between photoreceptor IS when viewed in longitudinally orientated samples (Fig. 4 B). Images from SBFSEM data showed these originated at the border between the outer nuclear (ONL) and the IS layers and run up to approximately half the length of the IS (Fig. S2, S3 A). When performing immunoEM labelling against actin (anti-β-actin in Fig. 4 C and phalloidin staining in Fig. S3 B), the staining was enriched within these projections as well as at membrane junctions at the proximal IS (white and black arrowheads respectively in Fig. 4 C). This was in contrast to the IS, where no cortical actin staining was detected at the lateral borders. A tomogram resolved the filamentous content running through the projection (Fig. 4 D), likely to be actin filaments. Labelling F-actin with phalloidin highlights the actin enriched projections within the IS layer and when tilting the 3D confocal data, a ‘honeycomb’ like pattern was observed reflecting the membrane junctions at the proximal IS (Fig. 4 E, Fig. S3 C and black arrow heads in Fig. 4 A-D). To determine the origin of the projections, an antibody against glutamine synthetase was used as it is a well-known Müller glial cell marker. By immunofluorescence there was no observed colocalization other than an overlapping region of enrich staining at the base of the projections (phalloidin staining) and within the glutamine synthetase channel (Fig. 4 F small panels). To investigate this further immunoEM labelling against glutamine synthetase was performed on retinal sections. This clearly showed that the projections emanated from the labelled Müller glial cells (Fig. 4 G and further staining in Fig. S3 D). In agreement with the IF staining the glutamine synthetase was absent from the projections in the immunoEM labelled sections, and only present within the cell body. By highlighting the projections within the SBFSEM data they could be traced to the cells that surrounded the photoreceptor rather than the photoreceptors themselves (Fig. 4 H), providing further evidence they are Müller glial cell derived.

**Fig. 4.**
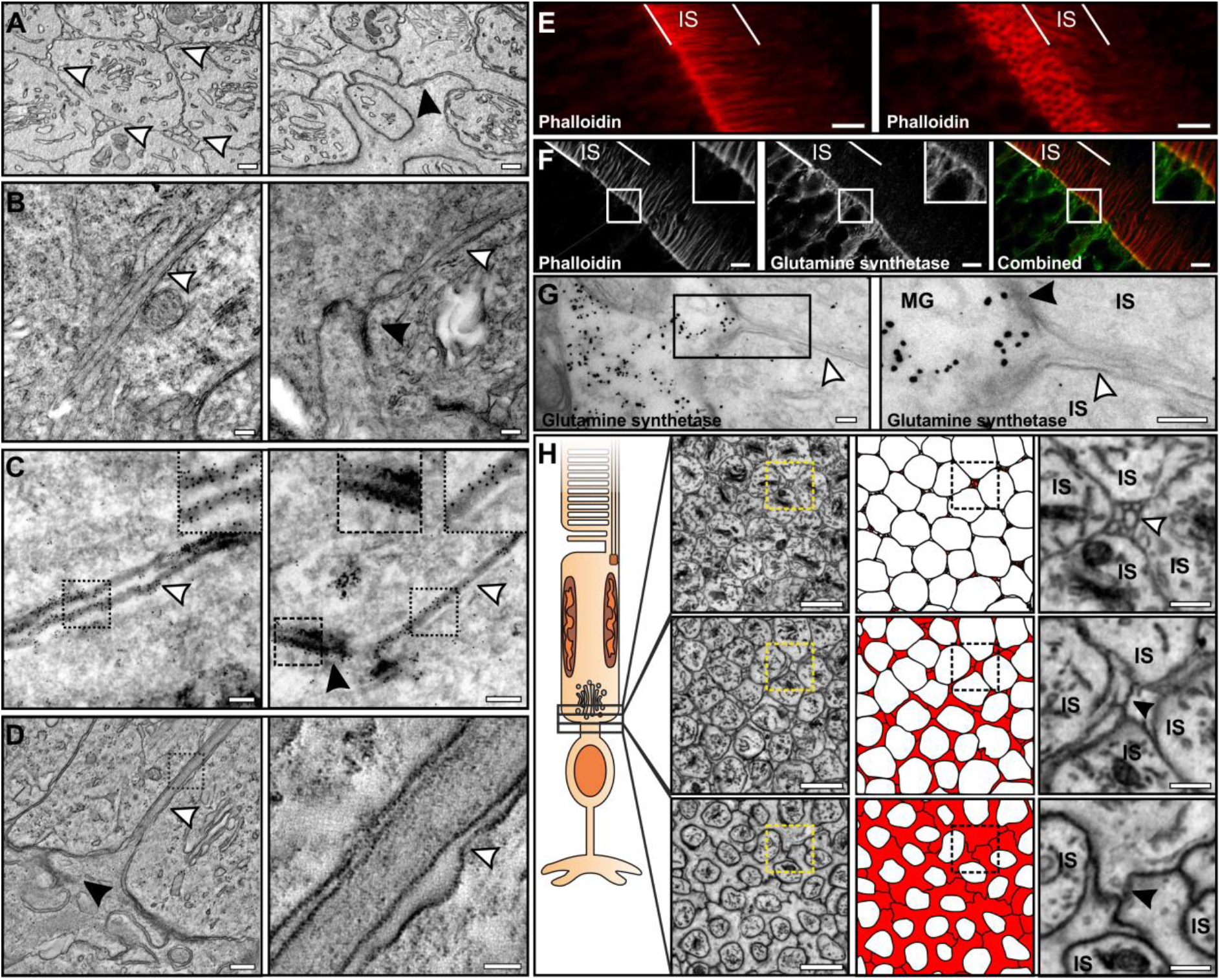
Projections that run between the ISs are actin rich and are Müller glial cell derived. (A) TEM images of cross sections at the proximal region of the IS. In the left panel projections run between the ISs (white arrowheads) and right panel is closer to the photoreceptor cell body, where the projections are no longer visible and junctions between the ISs are seen (black arrowhead). (B) Longitudinal images of the projections running between the ISs (indicated by white arrowheads, and black arrowhead shows the presence of junctions). (C) Immuno-EM labelling using an antibody against β-actin labelling indicates it is enriched within the projections (white arrowheads) and at junctions (black arrowhead). (D) A slice from a tomographic reconstruction, resolves the filamentous content of the projections. (E) Confocal stack of phalloidin stain retina, stains the projections and when tilting the stack (right panel) a ‘honeycomb’ pattern represents the actin enrichment at the junctions (shown by black arrowheads within the TEM images). (F) Immunofluorescent antibody labelling of phalloidin and the Müller glial marker glutamine synthetase that have little localisation (G) Immuno-EM labelling for glutamine synthetase shows the projections that lack labelling extend from the labelled Müller glial cells. (H) SBFSEM images show that the projections (coloured in red in the middle panel) originate from the Müller glial cells surrounding the photoreceptors in the outer nuclear layer. Scale: (A-D) 250nm, (E-F) 5um, (G) 200nm and (H) left handed panels 2μm and right handed panels 500nm.

### Mitochondria arrange against the IS PM between postnatal day 10 and 13

To determine the stage at which neighbouring photoreceptor IS mitochondrial align we examined mouse eyes pre- and post- full differentiation of the outer retina (before and after postnatal day 13 respectively) (17). At postnatal day 7 (P7) the immature photoreceptors (identified in the central retina by the presence of immature OSs, highlighted by OS in Fig. 5A) have a short IS consisting of scattered mitochondria, some in contact with the PM, as well as others positioned away from the PM (Fig. 5 A). At P7 we observed the presence of Müller glial processes, but there was no obvious alignment between mitochondria of neighbouring IS. By P10 a greater number of mitochondria were observed positioned against the PM with some still position centrally within the IS (Fig. 5 B). At this age some alignment between neighbouring IS mitochondria was observed. By P13 the mitochondrial positioning and alignment with neighbouring cell mitochondria reflected that of mature retina at P42 (Fig. 5 C, D white arrowheads).

**Fig. 5.**
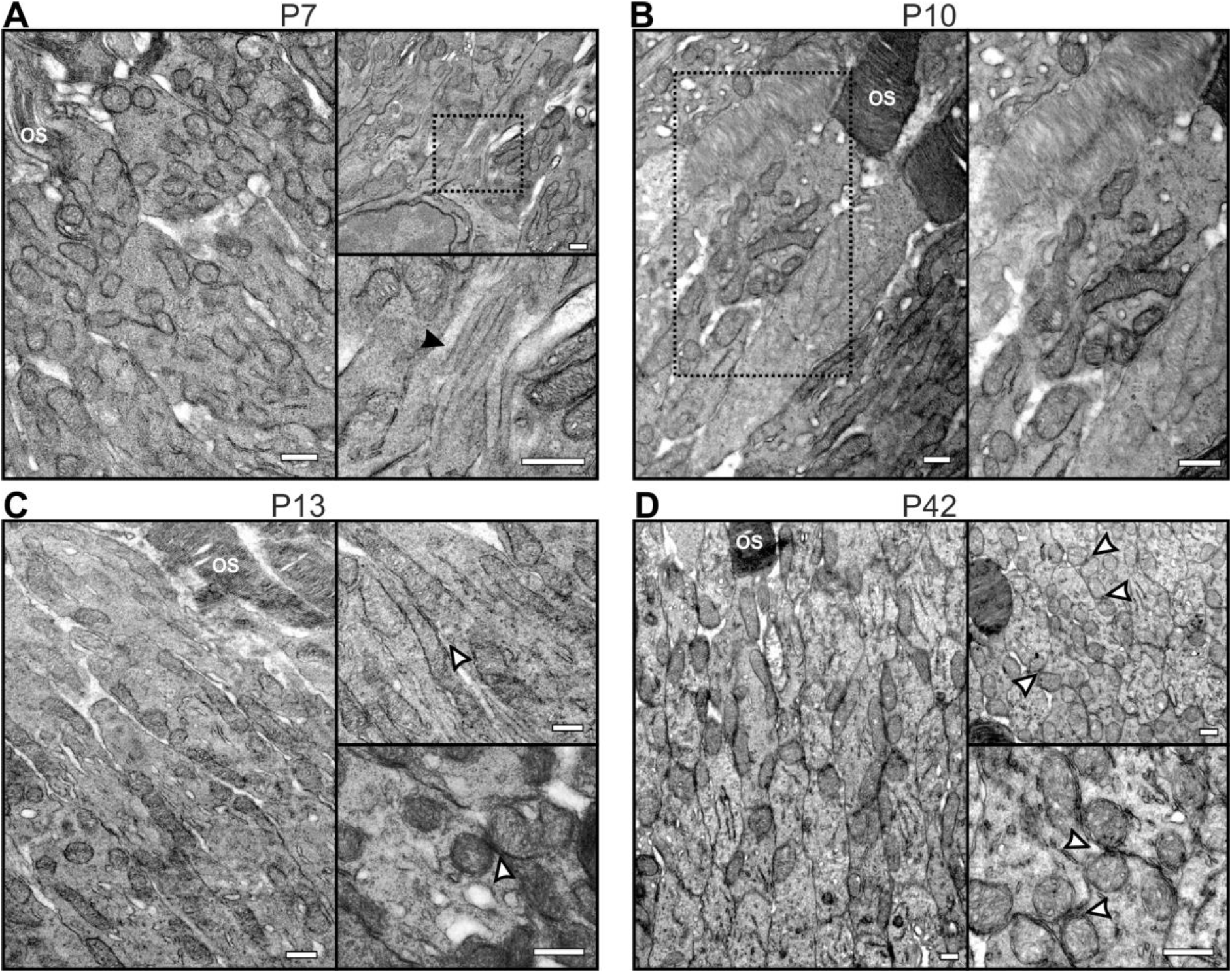
The mitochondria of neighbouring ISs are not aligned until postnatal day 13. (A-D) TEM images of ISs from mice of ages (A) P7, (B) P10, (C) P13 and (D) P42. The black arrowhead indicates the presence of Müller glial cell projections and white arrowheads alignment of neighbouring IS mitochondria. Scale: 500nm

### Heterozygous KO of Opa1 alters mitochondrial positing but does not affect cristae alignment

As OPA1 is known to play a role in mitochondrial fusion and cristae morphology (18) and specific deficits in visual electrophysiology (19), we examined the eyes of heterozygous KO mice by TEM. From longitudinally orientated retinal samples, the IS mitochondria of *Opa1* ^+/−^ mice presented an abnormal morphology and reduced alignment to neighbouring IS mitochondria when compared to *Opa1* ^+/+^ (Fig. 6A). Retinal cross sections at different depths of the IS showed some mitochondria in the *Opa1* ^+/−^ mice to be larger or abnormally shaped compared to those from *Opa1* ^+/+^ mice (Fig. 6B). The positioning of mitochondria within the IS and the mitochondrial diameter were quantified from photoreceptor cross-sections (Fig. 6 C-F). The percentage of IS containing mitochondria positioned away from the PM was higher in the *Opa1* ^+/−^ (38.32% ± 2.31% SE) compared to the *Opa1* ^+/+^ (13.25% ± 4.22% SE) mouse eyes (Fig. 6 C, E). Measurements of the shortest mitochondrial diameter indicated that the *Opa1* ^+/−^ mouse IS had larger mitochondria compared to the *Opa1* ^+/+^ mice (Fig. 6 D, F and Fig. S4 A). Tomographic reconstructions were generated to examine the mitochondrial cristae (Fig. 6 G). Within the tomographic slices, cristae openings were found to be aligned in mitochondria positioned against the PM in both the *Opa1* ^+/−^ and *Opa1* ^+/+^ IS (Fig. 6 G left-hand and central panel, as well Fig. S4 B). Large mitochondria positioned away from the PM in the *Opa1* ^+/−^ IS did not appear to have unusual or disordered cristae (Fig. S4 B).

**Fig. 6.**
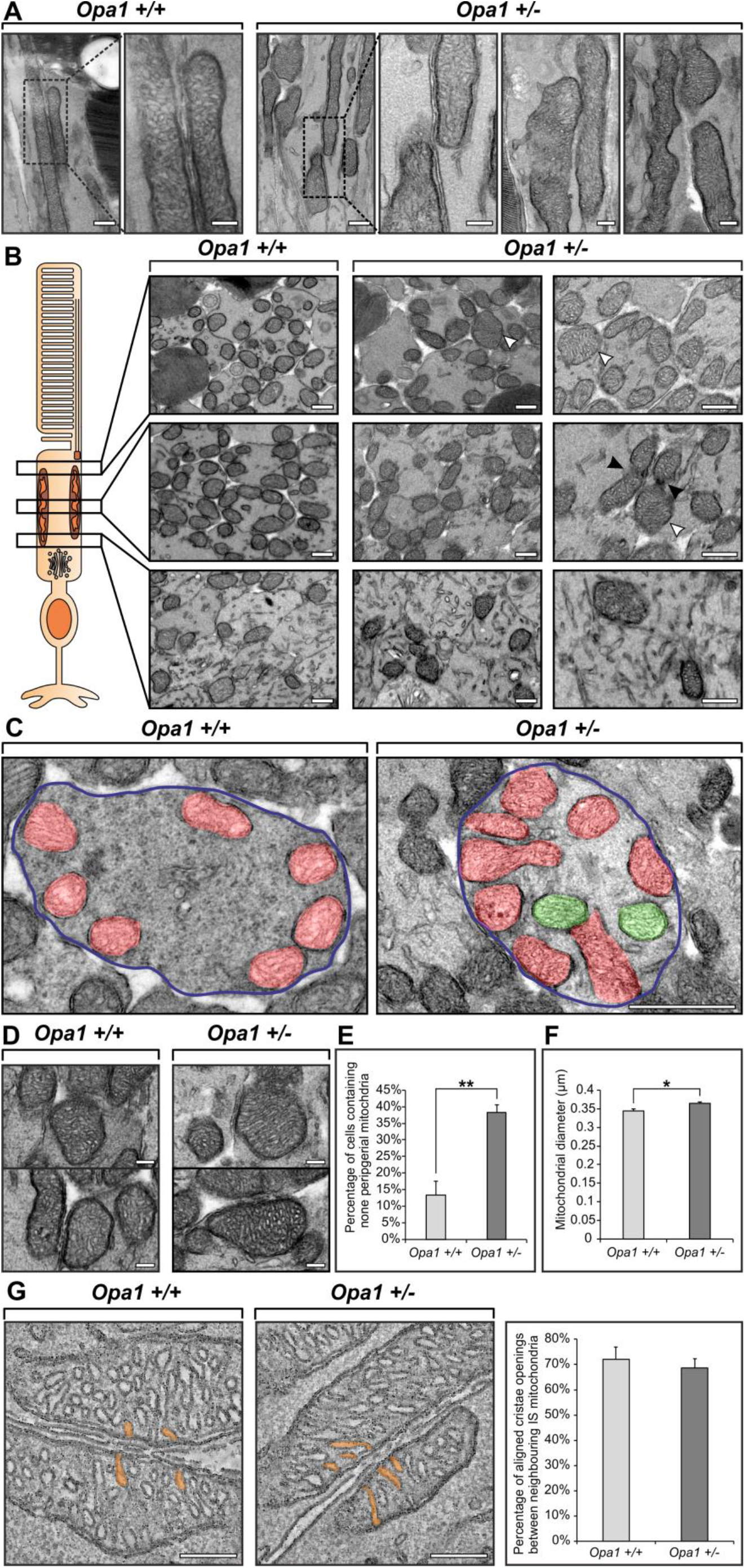
Heterozygous KO of *Opa1* leads to changes in the mitochondria morphology and positioning, but not the alignment of cristae openings. (A) TEM images of longitudinally orientated retina from *Opa1*+/+ and *Opa1*+/− mice. (B) Cross-sectioning orientated retina examined at different regions of the IS indicate enlarged mitochondria (white arrowheads) and evidence of a mitochondrial fusion defect (black arrowheads). (C) ISs cross sections in which mitochondria against the PM are false coloured in red and those away from the PM are in green. (D) Images showing larger mitochondria in the *Opa1*+/− mice ISs, but no obvious morphological defects of the cristae. (E) An increase in the number of ISs that have mitochondria positioned away from the PM in the *Opa1*+/− mice based on quantifying images similar to those shown in (C) (average = 3 eyes). (F) Measurements of mitochondrial show an increase in the average diameter in the *Opa1*+/− mice (average = 300 mitochondria from 3 eyes). (G) Tomographic slices shows the presence of cristae opening alignment in the both the *Opa1*+/+ and *Opa1*+/− mice, and measurement show no significant difference between the two mouse models (average = 3 tomograms). (E-G) Data are mean± SE, *P < 0.05, **P < 0.01 determined by Student t-test. Scale: (A) 200nm, (B) 500nm, (C) 1μm, (D & G) 200nm.

## Discussion

The high energy demands and importance of mitochondria for calcium storage in photoreceptor required for healthy vision are well established (1, 10). Yet the arrangement of mitochondria within photoreceptor IS has not been well studied. This has largely been hampered due to the lack of techniques to image through entire IS at the resolution that is achievable by TEM. The development of SBFSEM methodology allows serial imaging through large tissue volumes and is ideal for examining photoreceptor mitochondria. This combined with a range of electron microscopy techniques has allowed us to make a number of important and novel findings in regard to the highly specialised mitochondrial arrangement in the IS, shedding light on how this may help regulate energy and metabolite homeostasis across the mouse photoreceptor cell layer.

When examining through the depth of mouse photoreceptor IS, mitochondria remain in contact with the PM, and were seen to cluster together in pairs or triplets. The clustered mitochondria were found to be aligned to each other, running alongside most of the length of the IS, resulting in large mitochondrial surface areas facing each other. The distance between the mitochondria outer membrane and the PM was highly consistent, and we discovered this to be maintained by tethering between the membranes. This is the first time this kind of tethering has been shown in mammalian cells, which reflects the mitochondria–PM contact sites that have been studied in yeast (20). It has been suggested from previous studies that mitochondria are tethered to the PM in mammalian brain synapses (21). As these filamentous connections have been described as cytoskeletal anchors and run between mitochondria and adhering junctions over a distance greater than that of a contact site (>30 nm), they may not be linked directly to the PM and are a different type of tethering complex than the ~11nm tethers we have observed here (22–24). As there is no known mammalian homolog of the yeast mitochondria–PM tethering components Num1 and Mdm36, as well as a lack of a reliable *in vitro* photoreceptor model with fully differentiated OSs, we were unable to determine the constituents of the IS plasma tethers (20, 21). The close association of the mitochondria to the PM and the alignment to mitochondria from neighbouring photoreceptors implies there is communication and/or sharing of resources. Photoreceptors are highly sensitive to hypoxia and nutrient deprivation, but the pathway by which they can maintain the metabolite levels to meet their high energy demands is not fully understood (25). Furthermore, studies examining metabolic flux in the retina indicate the need for energy homeostasis across the photoreceptor cell layer and with the RPE to maintain retinal health and visual acuity (26–29). Therefore, the mitochondria arrangement described in this study may be an important evolutionary development for sharing the level of particular metabolites across the photoreceptor cell layer. It is possible the position of the mitochondria at the PM assists in directing light as there is evidence nocturnal mammals have a nuclear architecture consisting of heterochromatin localised in the centre that directs light up through the IS to the OS (30). Further work is required to test these hypotheses, using techniques to measure levels of metabolites as well as energy metabolism from photoreceptor cells in different mammalian models that have a clear mitochondrial disarrangement phenotype or disrupted mitochondrial tethering to the PM.

Tomographic reconstructions of IS mitochondria were used to resolve the fine cristae architecture. Single slices from the data as well as segmentation models showed the cristae opening showed a high degree of alignment between mitochondria of neighbouring cells. To determine if these were coincidental or exist in other cells types, RPE mitochondria were inspected. Mitochondria at the lateral RPE cell border were found to be in contact with the PM similar to what we found in IS, but most mitochondria from neighbouring cells were not found to be position side-by-side. Due to the conserved distance, we predict that there are tethers to the RPE lateral cell border, but in this study we were unable to clearly detect if they were present. When examining the cristae from the few neighbouring RPE cell mitochondria that were observed, the cristae showed little, if any, alignment when compared to what was observed in the IS. The alignment of the cristae opening further supports the notion of a mechanism for communication and/or regulate exchange of resources across PMs to mitochondria of neighbouring cells. Mitochondria within mouse cardiomyocytes and human skeletal muscle have been shown to form dense inter-mitochondrial junctions that have alignment/coordination of the cristae (31, 32). This is the first time cristae alignment has been observed between mitochondria from neighbouring cells. It is possible in photoreceptors that the mitochondria are polarised so that the cristae face the PM and the cristae opening are separated evenly, which gives the appearance of alignment. The tomography slices and model indicate this is unlikely, however, as the openings do not appear to be evenly distributed and a have a similar pattern of openings when compared to the opposing mitochondrial inner membranes. Furthermore, the shape of the cristae in some of the TEM images demonstrate it curving into position for alignment to a neighbouring mitochondrion. For the cristae to align across cells would require a coordinated complex at contact sites where the cristae opening are positioned. It is known at the cristae opening there is coordination and contacts between the inner and outer mitochondria membrane leading to the positioning of particular channels (33–35). At these sites on the outer mitochondrial membrane there would likely need to be further connections bridging to the PM, where there is another process that coordinates corresponding tethers within the neighbouring cells. For this to occur it is expected there would be either; (i) further tethers between cells similar to what we observed in tomography slices (Fig 2C), (ii) stimulation by released of metabolites/protein through PM channels or (iii) PM lipid enriched domains that are coordinated by the mitochondria to be positioned to line-up with cristae openings.

In the extracellular spaces between photoreceptors, often positioned close to the IS mitochondria, tube like projections were observed. In previously studies these have been proposed to be Müller glial processes, but there has been a lack of definitive evidence at the resolution of TEM to prove this to be the case (36, 37). To better characterise as well as determine the origin of these projections we used a combination of immuno-labelling and 3D electron microscopy techniques. We found that they were actin enriched, when staining for actin by immuno-EM, and while doing so we did not detect cortical actin at the lateral border of the photoreceptor IS. The latter finding correlates well with our mitochondria observations within the IS, as actin filaments would likely hinder the positioning and tethering of mitochondria to the PM. By staining for glutamine synthetase a well-known Müller glial cell marker, it was found to be absent from the projections, but labelled the cells that the projections emanated from. In addition to examining images through the depth of the retina close to the IS-ONL junction by SBFSEM, the projections do not originate from photoreceptors and we unequivocally show they are Müller glial cell derived. It has been proposed in retina there is an metabolic ecosystem, and in addition to the well-established metabolic transport between the RPE and photoreceptors (38–40), Müller glial cells are involved in shuttling of lactate as well as other metabolites (26). Due to the positioning of the Müller glial processes close to the IS mitochondria, they may exist to assist in transport of resources towards or away from the IS mitochondria. The actin filaments within the processes likely exists to support and stabilise the structure and other systems are involved in the transport of resources.

To determine the timepoint at which the mitochondria within the IS arrange against the PM aligning to neighbouring cell mitochondria, mouse eyes with developing photoreceptors were examined. At the youngest timepoint P7, the mitochondria appeared dispersed and gradually rearranged by P13, were the mitochondria where positioned in a similar arrangement to fully developed retina at P42. The outer limiting membrane layer that forms close to the proximal IS will have formed by P7, as Müller glial processes were observed, indicating their presence is independent to the mitochondrial rearrangement.

To determine the effect of reducing a known regulator of mitochondrial structure, Opa1, heterozygous *Opa1* KO mouse photoreceptors were examined. In the IS there was evidence of a fusion defect in some of mitochondria, but more strikingly the mitochondria were larger, and a greater proportion were position away from the PM in the *Opa1* ^+/−^ compared to the *Opa1* ^+/+^ mice. The cristae morphology and openings were found to be unaffected, and were aligned between neighbouring cell mitochondria in the *Opa1* ^+/−^ mice when examined in tomographic slices. Opa1 is most highly expressed in the retina and as heterozygous KO, the expression levels may have been adequate to provide normal cristae morphology, in combination with other factors that are regulating the cristae opening alignment (41). Due to the reduced proportion of mitochondria in contact with the PM, it makes the *Opa1* ^+/−^ a good model for future studies to determine the pathway that leads to the specialised IS mitochondrial arrangement.

This study sheds light on the importance on mitochondria in the IS and how their position and morphology have likely evolved to help fulfil the energy and storage demands across the photoreceptor cell layer. Further work is required to identify the tethers between the mitochondria and the PM as well as the factors regulating the cristae alignment. Furthermore, the positioning of mitochondrial and tether to the PM are likely to important in other cells types.

## Material and Methods

Mouse eyes were processed for SBFSEM, TEM, tomography, Immuno-EM and Immunofluorescence as described in detail in the SI. The eyes used were from mice that had been sacrificed by cervical dislocation in accordance with Home Office guidance rules under project licence 70/8101 and 30/3268. This was undertaken adhering to the Association for Research in Vision and Ophthalmology Statement for the Use of Animals in Ophthalmic and Vision Research. Heterozygous Opa1 KO mice were generate as previously described (42).

## Acknowledgements

We would like to thank Peter Munro for his help preparing the SBFSEM samples, Camilla Pilotti, Athina Dritsoula, Dimitrios Stampoulis and David Parfitt for providing wild-type mouse eye tissue and Astrid Limb for the glutamine synthetase antibody. This work was funded by grants from the Wellcome Trust (093445 to C.E.F and 205041 to M.E.C), Fight for Sight (1936UCL to M.C.S) and the Medical Research Council (G108523 and G0700949 to M.V).

## References

1. Kam JH, Jeffery G (2015) To unite or divide: mitochondrial dynamics in the murine outer retina that preceded age related photoreceptor loss. Oncotarget 6(29):26690–26701.

2. Wong-Riley M (2010) Energy metabolism of the visual system. Eye Brain 2:99–116.

3. Al-Enezi M, Al-Saleh H, Nasser M (2008) Mitochondrial Disorders with Significant Ophthalmic Manifestations. Middle East Afr J Ophthalmol 15(2):81–86.

4. Schrier SA, Falk MJ (2011) Mitochondrial Disorders and The Eye. Curr Opin Ophthalmol 22(5):325–331.

5. Giarmarco MM, Cleghorn WM, Sloat SR, Hurley JB, Brockerhoff SE (2017) Mitochondria Maintain Distinct Ca2+ Pools in Cone Photoreceptors. J Neurosci 37(8):2061–2072.

6. Križaj D (2012) Calcium Stores in Vertebrate Photoreceptors. Adv Exp Med Biol 740:873–889.

7. Fain GL, Matthews HR, Cornwall MC, Koutalos Y (2001) Adaptation in vertebrate photoreceptors. Physiol Rev 81(1):117–151.

8. Heidelberger R, Thoreson WB, Witkovsky P (2005) Synaptic transmission at retinal ribbon synapses. Prog Retin Eye Res 24(6):682–720.

9. Krizaj D, Copenhagen DR (2002) Calcium regulation in photoreceptors. Front Biosci J Virtual Libr 7:d2023–2044.

10. Szikra T, Križaj D (2007) Intracellular organelles and calcium homeostasis in rods and cones. Vis Neurosci 24(5):733–743.

11. Hudson G, et al. (2008) Mutation of OPA1 causes dominant optic atrophy with external ophthalmoplegia, ataxia, deafness and multiple mitochondrial DNA deletions: a novel disorder of mtDNA maintenance. Brain 131(2):329–337.

12. Alexander C, et al. (2000) OPA1, encoding a dynamin-related GTPase, is mutated in autosomal dominant optic atrophy linked to chromosome 3q28. Nat Genet 26(2):211.

13. Delettre C, et al. (2000) Nuclear gene OPA1, encoding a mitochondrial dynamin-related protein, is mutated in dominant optic atrophy. Nat Genet 26(2):207.

14. Meeusen S, et al. (2006) Mitochondrial Inner-Membrane Fusion and Crista Maintenance Requires the Dynamin-Related GTPase Mgm1. Cell 127(2):383–395.

15. Wang A-G, Fann M-J, Yu H-Y, Yen M-Y (2006) OPA1 expression in the human retina and optic nerve. Exp Eye Res 83(5):1171–1178.

16. Stone J, van Driel D, Valter K, Rees S, Provis J (2008) The locations of mitochondria in mammalian photoreceptors: Relation to retinal vasculature. Brain Res 1189:58–69.

17. Sharma RK, O’Leary TE, Fields CM, Johnson DA (2003) Development of the outer retina in the mouse. Dev Brain Res 145(1):93–105.

18. Frezza C, et al. (2006) OPA1 Controls Apoptotic Cristae Remodeling Independently from Mitochondrial Fusion. Cell 126(1):177–189.

19. Barnard AR, et al. (2011) Specific deficits in visual electrophysiology in a mouse model of dominant optic atrophy. Exp Eye Res 93(5):771–777.

20. Hammermeister M, Schödel K, Westermann B (2010) Mdm36 Is a Mitochondrial Fission-promoting Protein in Saccharomyces cerevisiae. Mol Biol Cell 21(14):2443–2452.

21. Westermann B (2015) The mitochondria-plasma membrane contact site. Curr Opin Cell Biol 35:1–6.

22. Perkins GA, et al. (2010) The Micro-Architecture of Mitochondria at Active Zones: Electron Tomography Reveals Novel Anchoring Scaffolds and Cristae Structured for High-Rate Metabolism. J Neurosci 30(3):1015–1026.

23. Prinz WA (2014) Bridging the gap: Membrane contact sites in signaling, metabolism, and organelle dynamics. J Cell Biol 205(6):759–769.

24. Spirou GA, Rowland KC, Berrebi AS (1998) Ultrastructure of neurons and large synaptic terminals in the lateral nucleus of the trapezoid body of the cat. J Comp Neurol 398(2):257–272.

25. Du J, Linton JD, Hurley JB (2015) Chapter Four - Probing Metabolism in the Intact Retina Using Stable Isotope Tracers. Methods in Enzymology, Metabolic Analysis Using Stable Isotopes., ed Metallo CM (Academic Press), pp 149–170.

26. Kanow MA, et al. (2017) Biochemical adaptations of the retina and retinal pigment epithelium support a metabolic ecosystem in the vertebrate eye. eLife 6:e28899.

27. Kooragayala K, et al. (2015) Quantification of Oxygen Consumption in Retina Ex Vivo Demonstrates Limited Reserve Capacity of Photoreceptor Mitochondria. Invest Ophthalmol Vis Sci 56(13):8428–8436.

28. Parapuram SK, et al. (2010) Distinct Signature of Altered Homeostasis in Aging Rod Photoreceptors: Implications for Retinal Diseases. PLOS ONE 5(11):e13885.

29. Wubben TJ, et al. (2017) Photoreceptor metabolic reprogramming provides survival advantage in acute stress while causing chronic degeneration. Sci Rep 7. doi:10.1038/s41598-017-18098-z.

30. Solovei I, et al. (2009) Nuclear Architecture of Rod Photoreceptor Cells Adapts to Vision in Mammalian Evolution. Cell 137(2):356–368.

31. Picard M, et al. (2015) Trans-mitochondrial coordination of cristae at regulated membrane junctions. Nat Commun 6:6259.

32. Vincent AE, et al. (2016) The Spectrum of Mitochondrial Ultrastructural Defects in Mitochondrial Myopathy. Sci Rep 6:30610.

33. Harner M, et al. (2011) The mitochondrial contact site complex, a determinant of mitochondrial architecture. EMBO J 30(21):4356–4370.

34. Hoppins S, et al. (2011) A mitochondrial-focused genetic interaction map reveals a scaffold-like complex required for inner membrane organization in mitochondria. J Cell Biol 195(2):323–340.

35. von der Malsburg K, et al. (2011) Dual Role of Mitofilin in Mitochondrial Membrane Organization and Protein Biogenesis. Dev Cell 21(4):694–707.

36. Uga S, Smelser GK (1973) Comparative Study of the Fine Structure of Retinal Müller Cells in Various Vertebrates. Invest Ophthalmol Vis Sci 12(6):434–448.

37. Woodford BarbaraJ, Blanks JanetC (1989) Localization of actin and tubulin in developing and adult mammalian photoreceptors. Cell Tissue Res 256(3). doi:10.1007/BF00225597.

38. Adijanto J, et al. (2014) The Retinal Pigment Epithelium Utilizes Fatty Acids for Ketogenesis IMPLICATIONS FOR METABOLIC COUPLING WITH THE OUTER RETINA. J Biol Chem 289(30):20570–20582.

39. Du J, et al. (2016) Reductive carboxylation is a major metabolic pathway in the retinal pigment epithelium. Proc Natl Acad Sci 113(51):14710–14715.

40. Senanayake P deS, et al. (2006) Glucose utilization by the retinal pigment epithelium: Evidence for rapid uptake and storage in glycogen, followed by glycogen utilization. Exp Eye Res 83(2):235–246.

41. Ham M, Han J, Osann K, Smith M, Kimonis V (2018) Meta-analysis of genotype-phenotype analysis of OPA1 mutations in autosomal dominant optic atrophy. Mitochondrion. doi:10.1016/j.mito.2018.07.006.

42. Davies VJ, et al. (2007) Opa1 deficiency in a mouse model of autosomal dominant optic atrophy impairs mitochondrial morphology, optic nerve structure and visual function. Hum Mol Genet 16(11):1307–1318.

43. Burgoyne T, et al. (2015) Rod disc renewal occurs by evagination of the ciliary plasma membrane that makes cadherin-based contacts with the inner segment. Proc Natl Acad Sci U S A 112(52):15922–15927.

44. Kremer JR, Mastronarde DN, McIntosh JR (1996) Computer visualization of three-dimensional image data using IMOD. J Struct Biol 116(1):71–76.

45. Slot JW, Geuze HJ, Gigengack S, Lienhard GE, James DE (1991) Immuno-localization of the insulin regulatable glucose transporter in brown adipose tissue of the rat. J Cell Biol 113(1):123–135.

46. Burgoyne T, Lane A, Laughlin WE, Cheetham ME, Futter CE (2018) Correlative light and immuno-electron microscopy of retinal tissue cryostat sections. PloS One 13(1):e0191048.

