## Supplementary Information for "Symmetric arrangement of mitochondria:plasma membrane contacts between adjacent photoreceptor cells regulated by Opa1"

**Supplementary Methods**

Transmission electron microscopy and tomography

Mouse eyes were fixed and embedded as described (43). ~100nm thick sections were cut and stained using lead citrate before acquiring images on a JEOL 1400+ TEM equipped with a Gatan Orius SC1000B charge-coupled device camera. For tomography 10-nm gold particle solution (fiducial marker) was used to stain the sections before tilting the stage from ±60° in 1.5° increments using the SerialEM software (University of Colorado Boulder). The images were processed and tomograms generated using the IMOD tomography package (44).

3view serial block face scanning electron microscopy

Eyes were fixed in 3% (vol/vol) glutaraldehyde and 1% (wt/vol) paraformaldehyde in 0.08 M sodium cacodylate buffer, pH 7.4 for 2 hrs at room temperature before incubating in the following solutions; 1% aqueous osmium tetroxide and 1.5% potassium ferrocyanide at 4°C for 1 hours, 1% aqueous thiocarbohydrazide at room temperature for 20 mins and 2% aqueous osmium tetroxide at room temperature for 30 minutes. They were sequentially en bloc stained 1% (wt/vol) aqueous uranyl acetate at 4°C overnight followed by Walton’s lead aspartate 30 mins at 60°C. The samples were dehydrated in an ethanol series followed by propylene oxide and infiltration in a mixture of propylene oxide and Durcupan ACM resin (1:1), before embedding in Durcupan ACM resin at 60° overnight. Blocks cut from the embedded specimens were mounted onto aluminum pins and coated with gold palladium. Using a Gatan 3View system (Gatan Inc, Abingdon, UK) and a Zeiss Sigma VP field emission scanning electron microscope (Zeiss, Cambridge, UK), images were acquired in between the sequential cutting away of 100nm thick section of the sample. The images were re-aligned using the StackReg plugin (EPFL) in ImageJ (NIH) and the images were modelled using the IMOD tomography package (44).

Cryo-immuno-electron microscopy

Mouse eyes were fixed in 4% (wt/vol) paraformaldehyde and 0.1% glutaraldehyde in 0.1 M phosphate buffer at pH 7.4 for 2hrs. The cornea and were lens removed before cutting the eye cup into small blocks and embedding them in 12% (wt/vol) gelatin, followed by infusion with 2.3 M sucrose solution at 4 °C overnight. 80nm sections were cut at −120 °C and collected in 1:1 mixture of 2.3 M sucrose/2% (wt/vol) methylcellulose, and labeling was performed as described previously (45). Labelling of actin was performed using an anti-β-actin antibody (Sigma) or phalloidin bound to biotin (Molecular Probes) and anti-biotin (Rockland), followed by protein-A-gold (CMC, University Medical Center Utrecht).

Immunofluorescence

The eyes were fixed in 4% (wt/vol) paraformaldehyde in PBS for 2 hrs before infusing with 30% (wt/vol) sucrose at 4°C before embedding in OCT compound and freezing using a bath of acetone cooled to -78°C. 5–20 um sections of the frozen samples were cut at -20°C using a cryostat. The sections were permeabilised using 0.1% (vol/vol) saponin in PBS for 30 mins at room temperature being antibody labelling in blocking solution consisting 0.01% (vol/vol) saponin in 1% (wt/vol) BSA in PBS for 2 hrs at room temperature. The sections were incubated in the following primary antibodies; glutamine synthetase (Novus Biologicals) and RETP1 (Abcam). Secondary antibodies bound to an Alexa Fluor dye (ThermoFisher Scientific) were applied to the sections for 1 hr at room temperature before mounting mounted with ProLong Gold antifade reagent (Life Technologies) that contained DAPI to counterstain the nuclei. Images were taken on a Leica SP8 confocal microscope.

Immuno-electron microscopy of cryostat sections

Mouse eyes were prepared following the methods described here (46). In brief cryostat sections were permeabilised using 0.05% (vol/vol) triton in PBS for 30 mins at room temperature and using a blocking solution containing 1% (wt/vol) BSA and 0.1% acetylated BSA in PBS, antibody labelling was performed using anti-glutamine synthetase (Novus Biologicals) for 2 hrs at room temperature. A secondary anti-rabbit bound to nano-gold (Nanoprobes) was applied in blocking solution for 2 hrs at room temperature before fixing the sections in 2% (vol/vol) glutaraldehyde, 2% (wt/vol) paraformaldehyde in 0.15 M sodium cacodylate buffer, pH 7.4 for 1 hr at room temperature. A gold enhance solution prepared in accordance with manufacturers specifications (Nanoprobes) and applied to the sections at 4°C for 10 mins. The sections were incubated in 1% (wt/vol) osmium tetroxide/1.5% (wt/vol) potassium ferrocyanide in distilled water for 1 hr at 4°C, before dehydrating in an ethanol series and embedding in epon at 60°C overnight. ~100nm thick sections were cut and images acquired on a JEOL 1400+ TEM with a Gatan Orius SC1000B charge-coupled device camera.

Image analysis

Electron microscopy and confocal images were analysed and measurements were made using ImageJ (NIH).

Fig. S1. Neighbouring photoreceptor mitochondrial alignment exists between cones and rods. The larger central cell (false coloured red) in the electron microscopy image is a cone that can be differentiated from rods due to its larger diameter and more electron lucent mitochondria. Alignment of mitochondria between cones and neighbouring rods indicated by the white arrowheads. Scale: 1µm

Fig. S2. Further examples of the mitochondrial alignment of neighbouring inner mitochondria as well as modelling of the projections seen in the extracellular space. (A-C) Segmentation of mitochondria from neighbouring inner segments and models generated showing them running side-by-side through the depth of the inner segment. (D-F) Segmentation of inner segment plasma membrane, mitochondria and projections. (G-L) The models generated from the segmentation show the projections are positioned close to mitochondria and run approximately half way up the inner segment. Scale: 1µm

Fig. S3. Further evidence that the projections are actin enriched and originate from Müller glial cells. (A) SBFSEM images showing the projections opening up into the Müller glial cells surrounding the photoreceptors. (B) Immuno-EM of thawed frozen sections using phalloidin conjugated to biotin followed by anti-biotin labelling and 10nm protein-A-gold. The gold labelling localises to the projections indicating actin enrichment. (C) Immunofluorescence using an antibody against rhodopsin (green) as well as phalloidin (red) and DAPI (blue) staining. (D) Immuno-EM labelling of cryostat sections using antibody against the Müller glial cell marker glutamine synthetase. Scale: (A) 500nm, (B) 200nm, (C) 5µm, (D) left-handed panels 500nm and right-handed panels 250nm.

Fig. S4. Alternative representation of the mitochondria diameters measured and further tomograms showing evidence of cristae alignment in the Opa1 heterozygous knockout. (A) The distribution of 300 mitochondria diameters measured from the photoreceptor inner segments of 3 eyes from both *Opa1*+/+ and *Opa1+*/- mice. (B) Further tomographic slices showing the existence of aligned cristae opening in the *Opa1*+/+ and *Opa1+*/- mice and the far right-side panel show large mitochondria positioned away from the plasma membrane in the *Opa1+*/- mice have normal looking cristae. Scale: (B) 200nm.

Movie S1. Tomogram of mouse rod photoreceptor IS mitochondria from Fig. 2.

Movie S2. Further tomogram of mouse rod photoreceptor inner segment mitochondria from Fig. 2.

Movie S3. Model of mitochondrial inner and outer membranes and plasma membranes from two neighbouring photoreceptor inner segments.
