## Supplementary figures and images for "Symmetric arrangement of mitochondria:plasma membrane contacts between adjacent photoreceptor cells regulated by Opa1"

### Supplementary Figure 1

Supplementary Figure 1

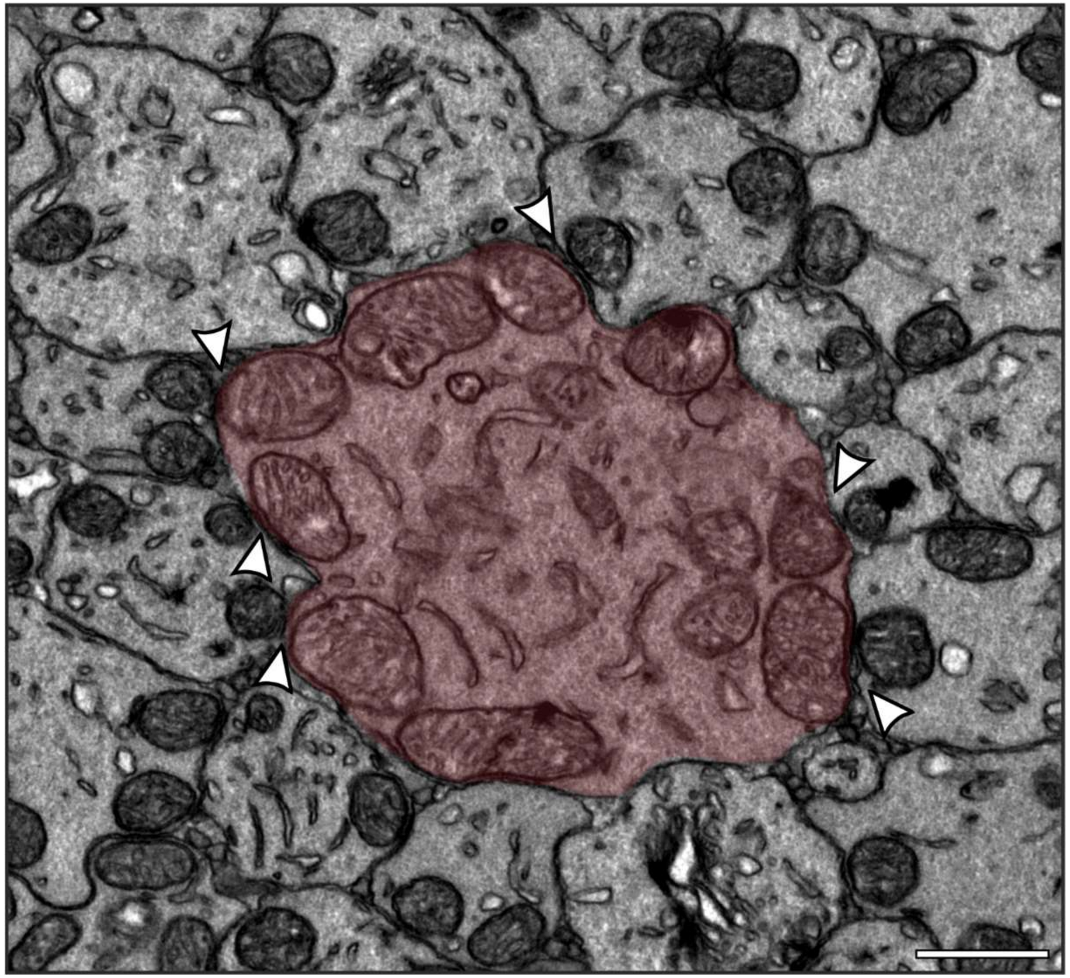

### Supplementary Figure 2

#### Supplementary Figure 2

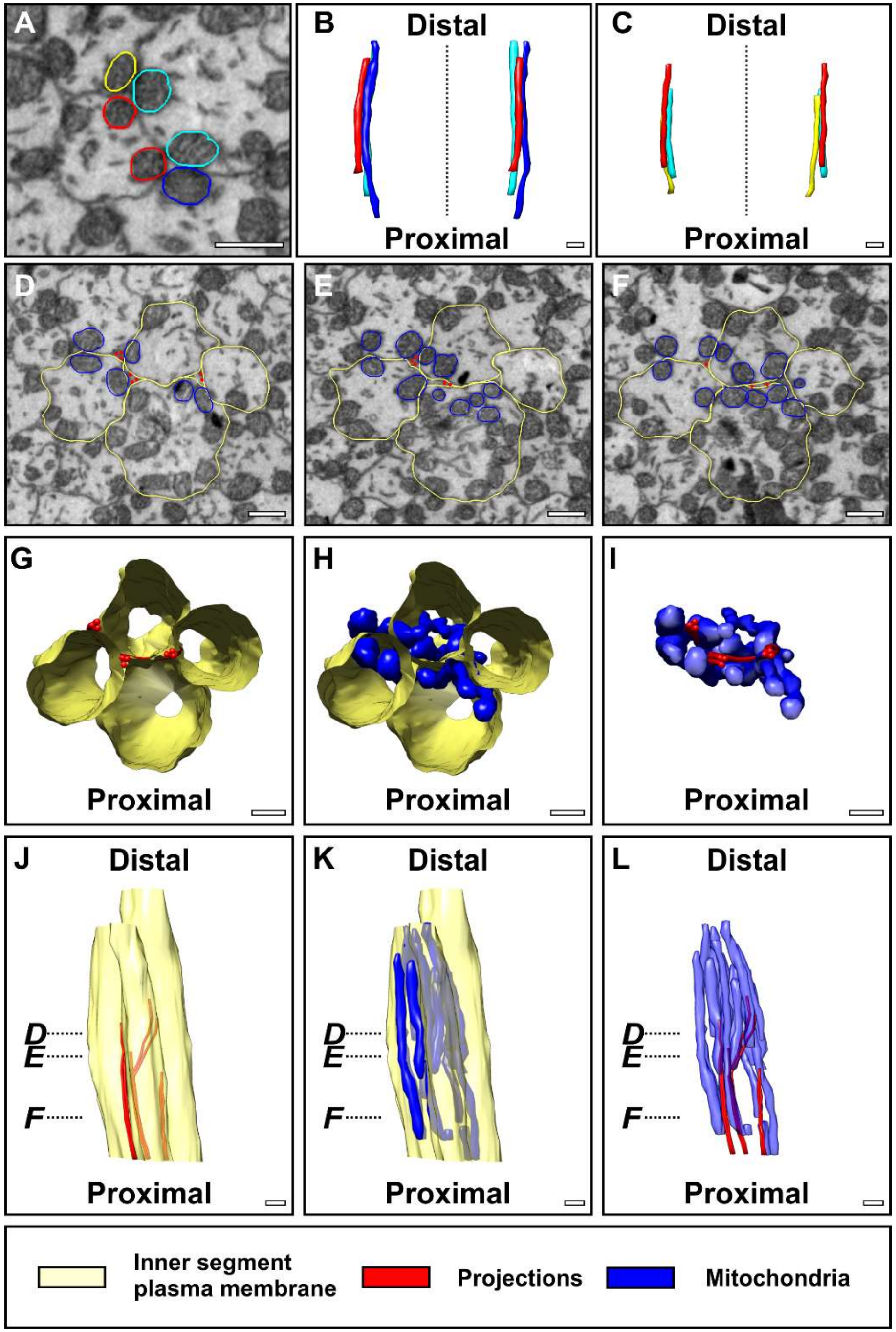

### Supplementary Figure 3

Supplementary Figure 3

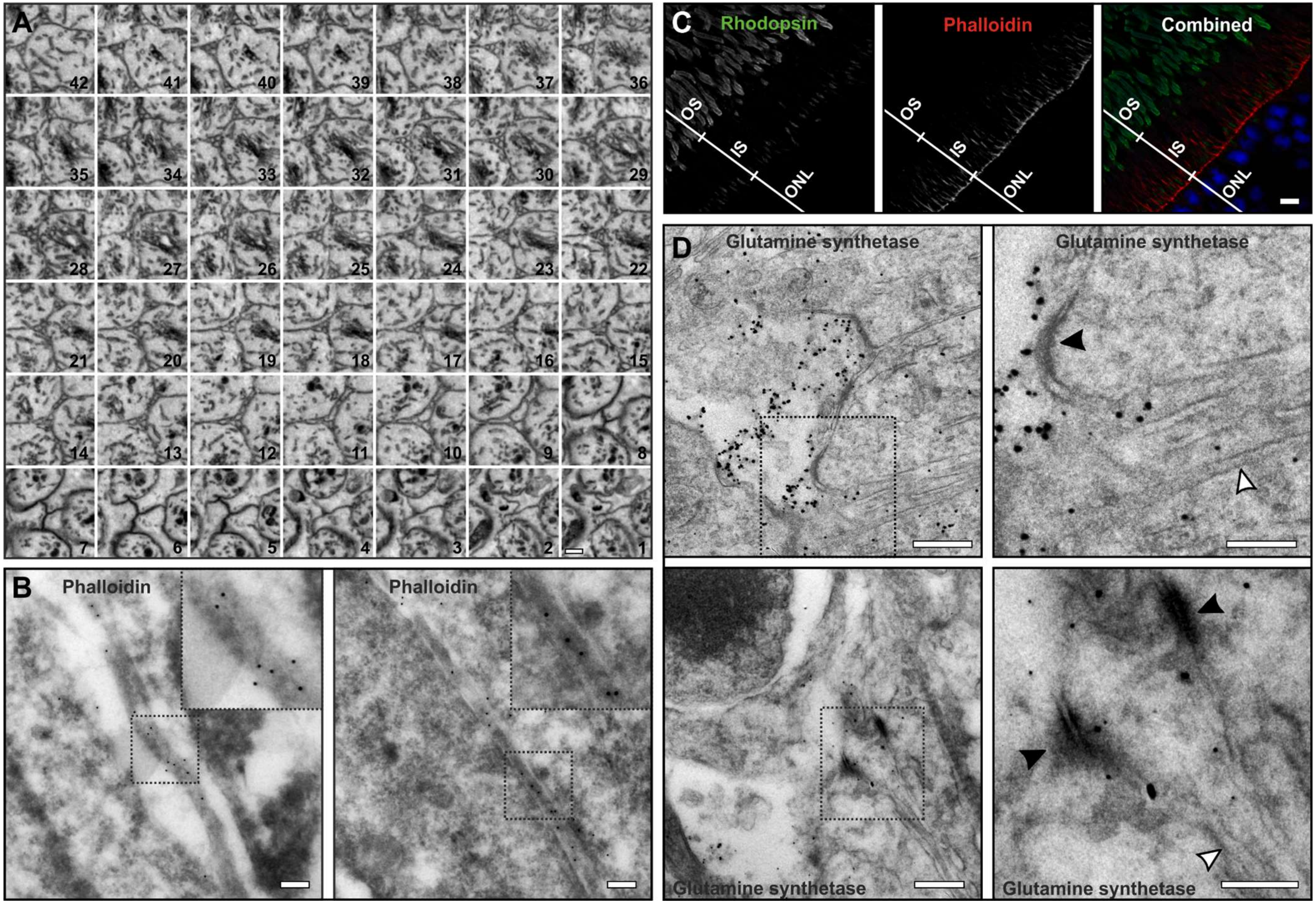

### Supplementary Figure 4

#### Supplementary Figure 4

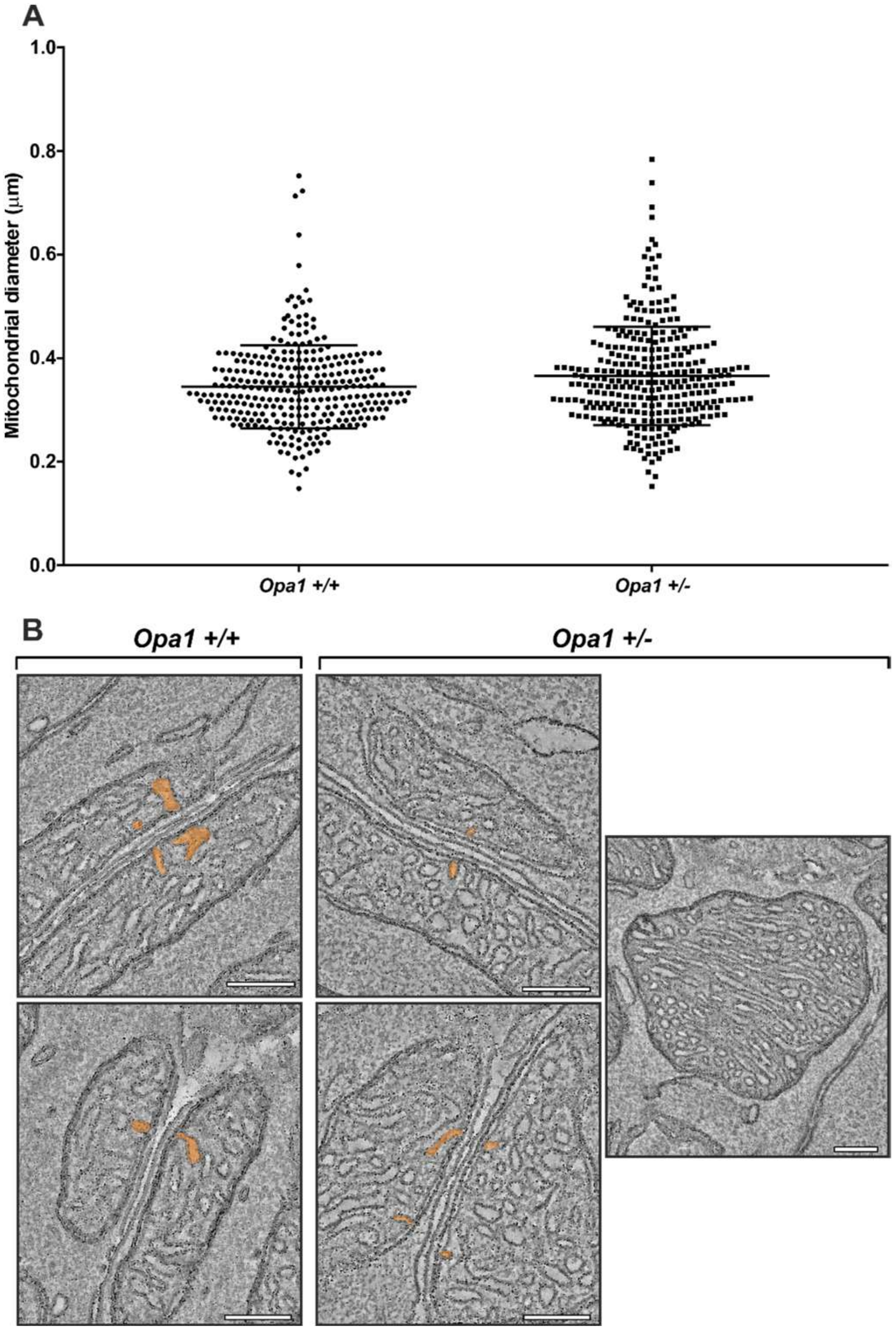
